# Cortical Representation of Movement Across the Developmental Transition to Continuous Neural Activity

**DOI:** 10.1101/2023.01.22.525085

**Authors:** Ryan Glanz, Greta Sokoloff, Mark S. Blumberg

## Abstract

Primary motor cortex (M1) exhibits a protracted period of development that includes the establishment of a somatosensory map long before motor outflow emerges. In rats, the sensory representation is established by postnatal day (P) 8 when cortical activity is still “discontinuous.” Here, we ask how the representation survives the sudden transition to noisy “continuous” activity at P12. Using neural decoding to predict forelimb movements based solely on M1 activity, we show that a linear decoder is sufficient to predict limb movements at P8, but not at P12; in contrast, a nonlinear decoder effectively predicts limb movements at P12. The change in decoder performance at P12 reflects an increase in both the complexity and uniqueness of kinematic information available in M1. We next show that the representation at P12 is more susceptible to the deleterious effects of “lesioning” inputs and to “transplanting” M1’s encoding scheme from one pup to another. We conclude that the emergence of continuous cortical activity signals the developmental onset in M1 of more complex, informationally sparse, and individualized sensory representations.

## Introduction

Perhaps the most striking feature of infant cortical activity is its discontinuity: Periods of silence are punctuated by bursts of population-level activity [1, 2]. This early phase of discontinuity ends with the sudden and dramatic onset of continuous cortical activity. Whereas in humans this transition occurs around the time of birth [3], in infant rats and mice it occurs at the end of the second postnatal week [4-8]. In fact, the onset of continuous cortical activity is but one aspect of a more global reorganization of brain dynamics that includes a shift in GABAergic functioning [9, 10], the increase and diversification of inhibitory interneurons [11-14], accelerated myelin deposition [15], and the onset of brain rhythms such as delta [16-18] and theta [19].

In primary motor cortex (M1), continuous activity emerges in rats between postnatal days 8 and P12 [6]. Despite its name, M1 at these ages does not produce movement, but instead functions exclusively as a somatosensory structure several weeks before the emergence of motor outflow [20-22]. Recently, we showed that neurons within M1’s nascent somatosensory map are tuned to limb kinematics, as is the case for M1’s adult motor map [23, 24]. Specifically, we found precise sensory tuning to movement amplitude at P8—especially for the limb twitches that occur during active (REM) sleep [6]. However, upon the emergence of continuous activity at P12, this tuning disappeared.

Was M1’s kinematic tuning truly lost, or was it only obscured within the noise of continuous activity? Here, we address this and related questions using a computational technique—*neural decoding*—that allows us to predict the timing and amplitude of forelimb movements based solely on M1 activity (see [25]). Our findings demonstrate that M1’s processing of sensory information is intact but substantially transformed after this fundamental transition in cortical dynamics.

## Results

We performed neural decoding using a previously published dataset from P8 and P12 rats (n = 8 at each age; [6]). This dataset consists of one-hour recordings of M1 unit activity and video-based records of 3-D forelimb displacement. For all decoding procedures, M1 unit activity and forelimb displacement served as the predictor and target variables, respectively. Each recording was divided into training (36 min; 60%) and testing (9 min; 15%) datasets; a validation dataset (15 min; 25%) was held back until the final model parameters were established. All analyses were performed on the validation dataset to ensure an unbiased assessment of decoder performance [25].

### Movement encoding is obscured by continuous activity at P12

Discontinuous M1 activity presents a strong contrast between periods of movement (when reafference triggers neural activity) and rest (when there is little to no activity). Moreover, at P8, we previously found that forelimb twitches trigger rate-coded M1 responses that correlate with twitch amplitude [6]. Accordingly, we predicted that a linear decoder would accurately predict the temporal and spatial properties of forelimb movements at P8. In contrast, because continuous activity at P12 occludes the temporal and spatial relationships between M1 activity and forelimb movements, we predicted that a linear decoder would no longer predict forelimb movements at that age. Indeed, by comparing actual with predicted limb displacement using a linear decoder (**Figure 1A**, blue lines), we confirmed both predictions: A linear decoder is sufficient to predict forelimb movements at P8, but not at P12.

**Figure 1.**
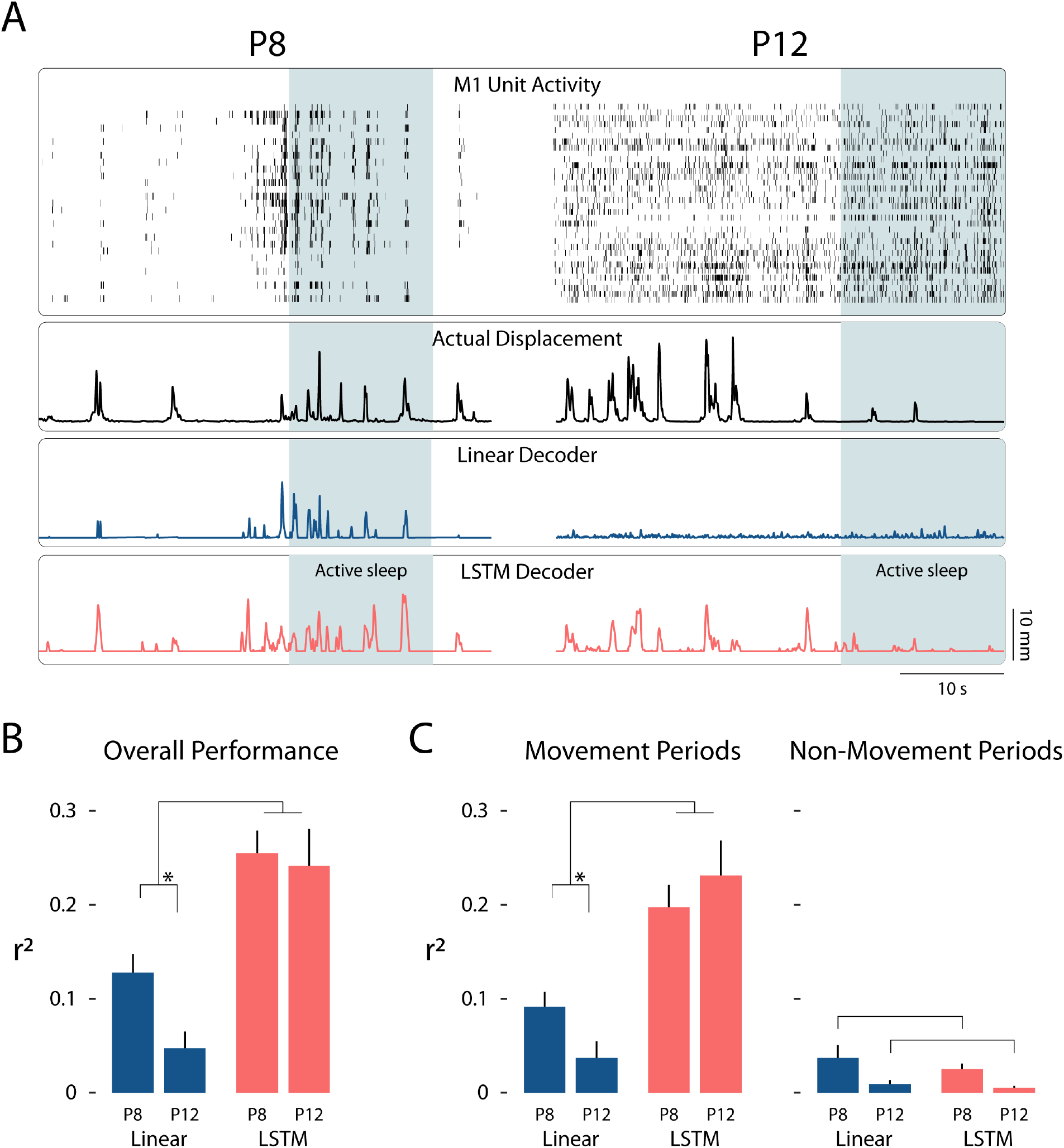
Neural decoding in M1 predicts forelimb movements across the transition to continuous cortical activity. **(A)** Representative data from a P8 (left) and P12 (right) rat. From top to bottom: M1 unit activity where each row denotes an individual unit and each vertical tick denotes an action potential; actual forelimb displacement (black lines) in mm, representing the Euclidean distance traveled by the forelimb in 3-dimensional space; forelimb displacement as predicted by a linear decoder (blue lines); forelimb displacement as predicted by an LSTM nonlinear decoder (orange lines). Shaded blue regions represent periods of active sleep. (**B**) Mean (+SEM) overall performance of the linear (blue bars) and LSTM (orange bars) decoders for P8 and P12 rats, as measured by *r*^2^ (the proportion of variance in forelimb displacement explained by the decoder). Brackets denote a significant interaction between age and decoder (p < .05). Asterisk denotes significantly better performance of the linear decoder at P8 compared to P12 (p < .05). (**C**) Left: Same as in (B), but for periods of forelimb movement. Right: Same as in (B), but for non-movement periods; brackets denote that both decoders performed significantly better at P8 than at P12 (p < .05).

Although we observed a loss of decoding accuracy at P12, M1 activity is nonetheless highly correlated with limb kinematics in adults [23, 24]. Accordingly, we hypothesized that M1 activity continues to represent movement kinematics after the emergence of continuous activity, but that this representation is too complex to be revealed using a linear decoder. We tested this hypothesis using a nonlinear decoder (long short-term memory decoder, LSTM; [26]) and found that it successfully decoded forelimb movements at P12, as well as P8 (**Figure 1A**, orange lines).

To quantify decoder performance, we computed the proportion of variance explained (*r*^2^) between the actual and predicted forelimb displacement for each decoder (**Figure 1B**). This metric represents the amount of temporal and spatial kinematic information available in M1 immediately after a forelimb movement. There was a significant interaction between pup age and the decoder used (F(1, 14) = 19.15, p < .001, adj. 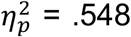; **Figure 1B**), indicating (1) that the linear decoder performed significantly better at P8 than P12 (F(1, 14) = 10.08, p = .007, adj.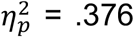), and (2) that the nonlinear decoder significantly outperformed the linear decoder at both ages. Thus, M1 activity indeed preserves kinematic information after continuous activity emerges at P12, though this relationship is only observable using a nonlinear decoder.

### State-dependent encoding of movement-related information

To confirm that decoder performance was driven by M1 activity specifically during periods of limb movement, the recordings were split into movement and non-movement periods, as described previously [6]. As expected, successful decoder performance was attributable to movement periods (F(1, 14) = 7.35, p = .017, adj.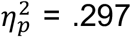; **Figure 1C, left**) and not to non-movement periods (F(1, 14) = 0.06, p = .817, adj.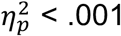; **Figure 1C, right**). Although there was a main effect of age on decoder performance during non-movement periods (F(1, 14) = 9.15, p = .009, adj. 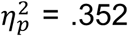), this effect was negligible compared with the performance of the decoder during movement periods.

The last analysis did not distinguish between movements during sleep (i.e., twitches) and wake. Thus, we next compared decoding accuracy for twitches and wake movements, focusing on a 2-s window centered on each forelimb movement (**Figure 2A**). For twitches, a main effect of age indicated that decoding performance was significantly better at P8 than at P12, regardless of whether a linear or non-linear decoder was used (F(1, 3808) = 534.25, p < .001, adj.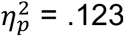; **Figure 2B, left**). Unexpectedly, the nonlinear decoder’s performance for twitches did not improve on the linear decoder’s performance at either age (F(1, 3808) = 1.67, p = .197, adj.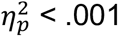). This finding suggests that M1 activity no longer represents twitch kinematics after continuous activity emerges (although reafference from twitches continues to trigger M1 activity through at least P20; [27]).

**Figure 2.**
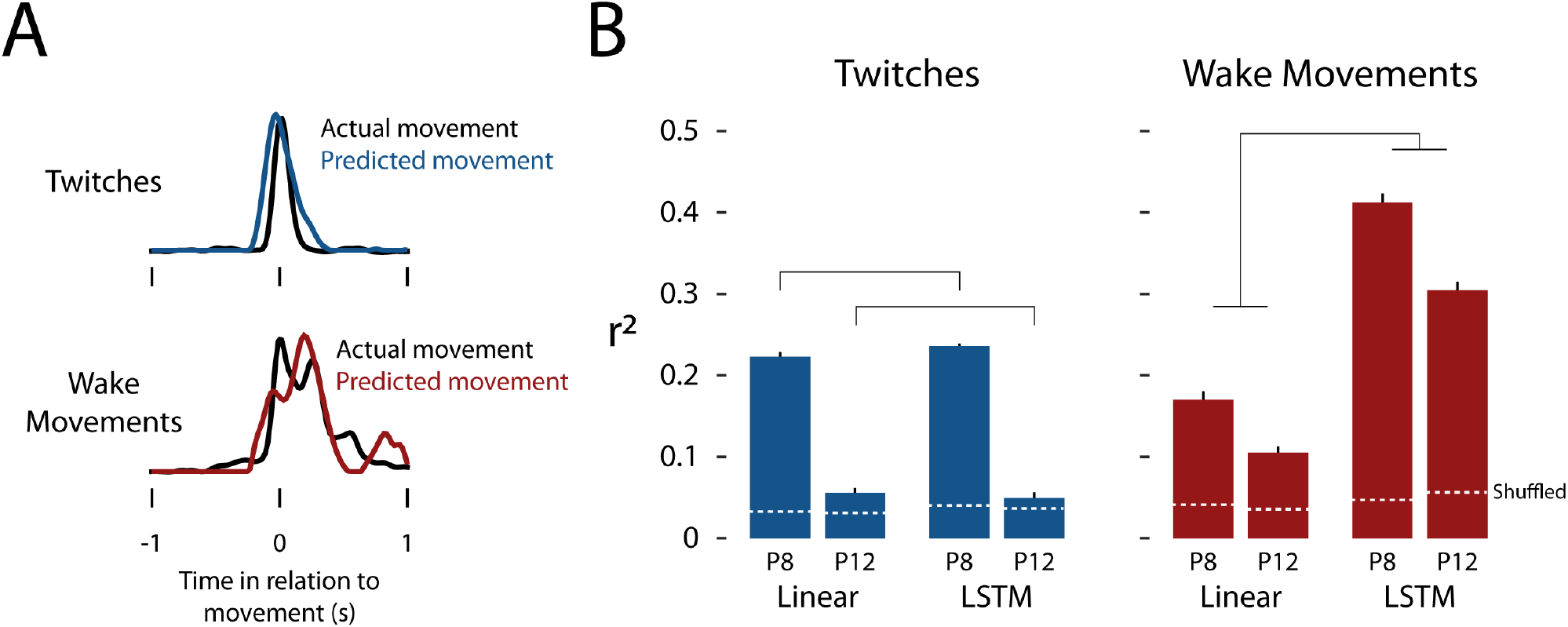
Decoding performance for twitches and wake movements varies by age. (**A**) Forelimb displacement for actual movements (black lines) compared to a predicted twitch (blue line, top) and a predicted wake movement (red line, bottom). A single representative twitch and wake movement is shown. (**B**) Mean (+SEM) decoder performance (*r*^2^) for twitches (left; blue bars) and wake movements (right; red bars). For each type of movement, the linear decoder is compared to the LSTM decoder across P8 and P12 rats. For twitches, brackets denote that both decoders performed significantly better at P8 than at P12 (p < .05). For wake movements, brackets denote that the LSTM decoder performed significantly better than the linear decoder at both ages (p < .05). Horizontal dashed white lines indicate chance performance using a shuffling procedure.

In contrast with twitches, for wake movements the nonlinear decoder performed significantly better than the linear decoder at P8 and P12 (F(1, 1340) = 240.91, p < .001, adj.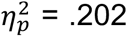; **Figure 2B, right**). This finding was particularly surprising at P8 because reafference from wake movements is blunted at this age [20, 21]. Thus, wake movements appear to be represented in a nonlinear fashion even as early as P8, and this representation survives the transition to continuous activity at P12. For both decoders, there was a small but statistically significant decrease in performance at P12 compared with P8 (F(1, 1340) = 18.24, p < .001, adj.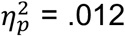), suggesting that the emergence of continuous activity modestly interferes with M1’s representation of wake movements. Together, these results indicate that although M1’s representation of twitch kinematics diminishes between P8 and P12, its representation of wake-movement kinematics is robust at both ages.

### Continuous activity increases the uniqueness of M1 information

The presence of decorrelated, continuous M1 activity at P12 represents a transition from a “dense” (i.e., redundant) encoding of information to a more energy- and information-efficient “sparse” code [28]. Accordingly, we predicted that removal of redundant neural input at P8 would not degrade decoding performance as much as removal of sparse neural input at P12.

To test this hypothesis, two “lesion” experiments were performed. In the first “unit lesion” experiment, a varying percentage of M1 units (up to 90%) recorded from a given pup was randomly discarded from the decoding process (**Figure 3A, top**). The resulting decoding performance was then compared with the decoding performance of the “intact” model (i.e., 0% of units removed). (Only the nonlinear decoder was used for these experiments.) As predicted, although unit lesions degraded decoder performance at both ages, the loss was significantly greater at P12 than at P8 (F(1.50, 20.94) = 18.60, p < .001, adj.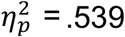; **Figure 3A, bottom**). For example, lesioning 50% of units led to a decoder performance of 70.4 ± 3.9% at P8, compared with 41.9 ± 4.4% at P12. Such differences in decoder performance were statistically significant across the range of unit lesions (all ps ≤ .005).

**Figure 3.**
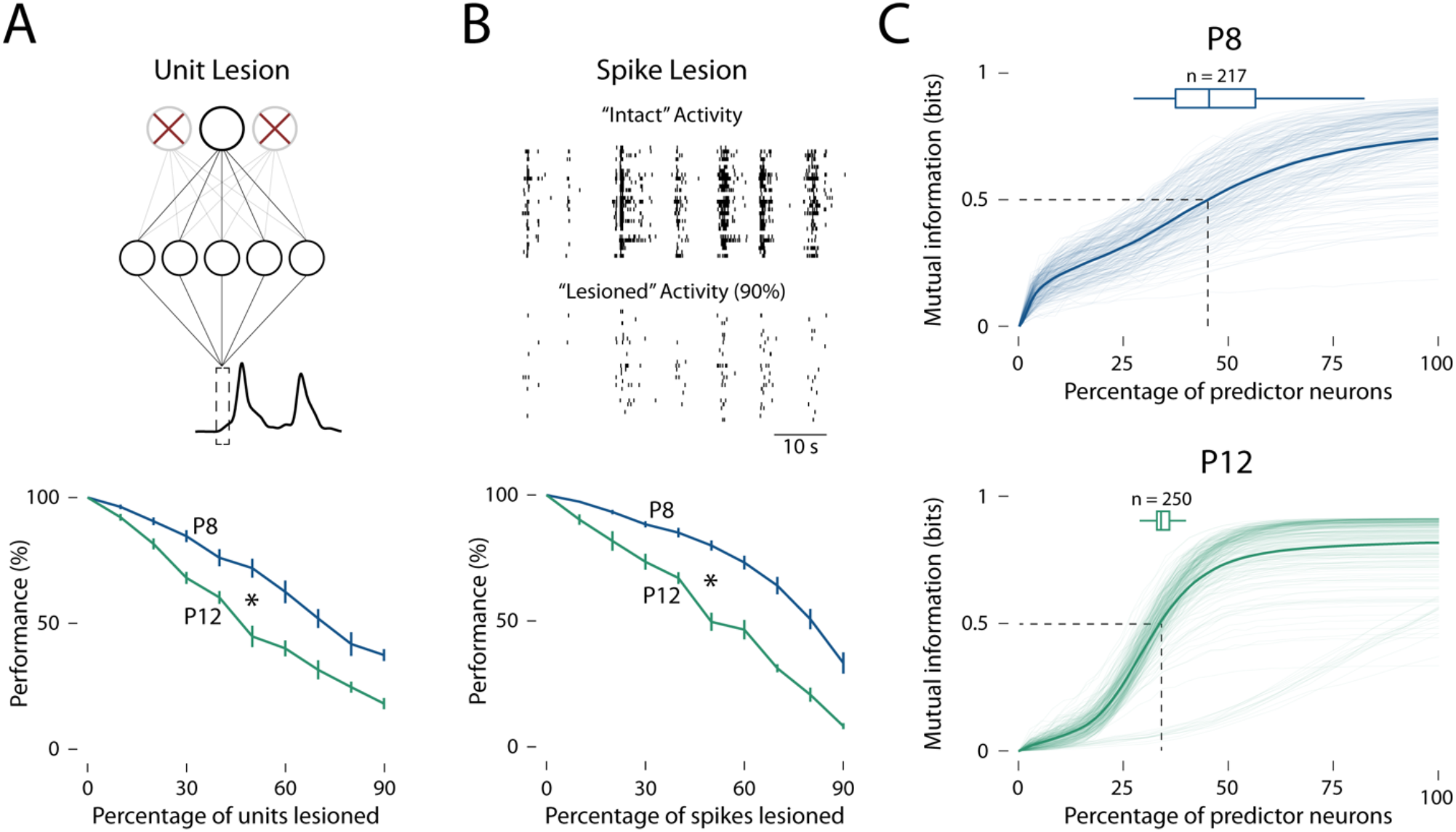
M1 activity contains more unique information at P12 than at P8. (**A**) Top: Representation of the “unit lesion” experiment. The top row of nodes represents the individual M1 units input to the LSTM decoder. Red X’s denote units that were “lesioned” (i.e., removed from the decoding process). The second row of nodes represents the hidden layer of the LSTM decoder. The black trace at the bottom represents forelimb displacement as predicted by the LSTM decoder. Bottom: Mean (±SEM) decoder performance (measured as the percentage change in the “lesioned” *r*^2^ relative to the “intact” *r*^2^) for P8 (blue line) and P12 (green line) rats as a function of the percentage of units lesioned. Asterisk denotes significantly worse decoder performance at P12 than at P8 across the range of units lesioned (p < .05). (**B**) Top: Representation of the “spike lesion” experiment. “Intact” activity represents baseline M1 activity, where each row represents an individual unit and each vertical tick denotes an action potential. “Lesioned” activity represents M1 activity after 90% of individual spikes were removed from the decoding process. Bottom: Same as in (A), but for the spike lesion experiment. Asterisk denotes significantly worse decoder performance at P12 than at P8 across the range of spikes lesioned (p < .05). (**c**) For P8 (top, blue) and P12 (bottom, green) rats, mutual information (in bits) is shown as a function of increasing size of the predictor subset (expressed as a percentage of the maximum size). Solid lines represent the mean increase in mutual information in the target unit; translucent lines represent individual target units (P8: n = 217 units; P12: n = 250 units). Dashed lines show the mean percentage of predictor units required to achieve 0.5 bits (half the theoretical maximum); horizontal boxplots show the percentage of predictor units required to achieve 0.5 bits for individual target units.

The second “spike lesion” experiment was performed in a similar fashion. Here, a varying percentage of spikes (i.e., action potentials) across all M1 units (up to 90%) was randomly discarded from the decoding process (**Figure 3B, top**). As predicted, spike lesions resulted in significantly greater declines in decoder performance at P12 than at P8 (F(5.05, 70.63) = 8.41, p < .001, adj.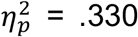; **Figure 3B, bottom**). For example, lesioning 50% of spikes led to a decoder performance of 79.5 ± 1.8% at P8, compared with 51.0 ± 3.2% at P12. Again, such differences in decoder performance were statistically significant across the range of spikes removed (all ps ≤ .03).That both lesions decreased decoder performance more at P12 than P8 implies that M1 activity contains more *unique* information about limb kinematics at the older age. To test this implication, we measured the mutual information between a target M1 unit and a randomly selected subset of units [29]; the size of the subset varied from 1 additional unit to all available units for a given pup. Here, mutual information refers to the predictability of the target unit’s response (to a movement) when the responses of the subset are known. By definition, mutual information *only* increases when the subset contains non-redundant (i.e., unique) information about the target unit [30]. We found that adding units increased mutual information significantly more quickly at P12 (38.7 ± 1.0% units to reach 0.5 bits) than at P8 (50.9 ± 1.2% units to reach 0.5 bits; t(436.01) = 7.64; p ≤ .001, adj. η_*p*_^2^= .116; **Figure 3C**; note that 1 bit is the theoretical maximum value). Together, these results support the notion that the onset of continuous activity at P12 increases the uniqueness of information available in M1.

### M1’s encoding scheme is interchangeable at P8, but not at P12

The finding that M1’s representation of movement at P8, but not at P12, is so redundant as to be robust to the removal of neural activity raises an intriguing hypothesis: Namely, that discontinuous activity is not only redundant, but perhaps is also generic to the point of being interchangeable between individual pups. Such a property of M1 activity could reflect a gross encoding scheme that is present during this period of development when somatotopic relations among M1 units and forelimb muscles are still being established. Accordingly, we predicted that after “transplanting” the encoding scheme of one pup into that of another, decoding performance would remain high at P8, but would be degraded at P12 (**Figure 4A**).

**Figure 4.**
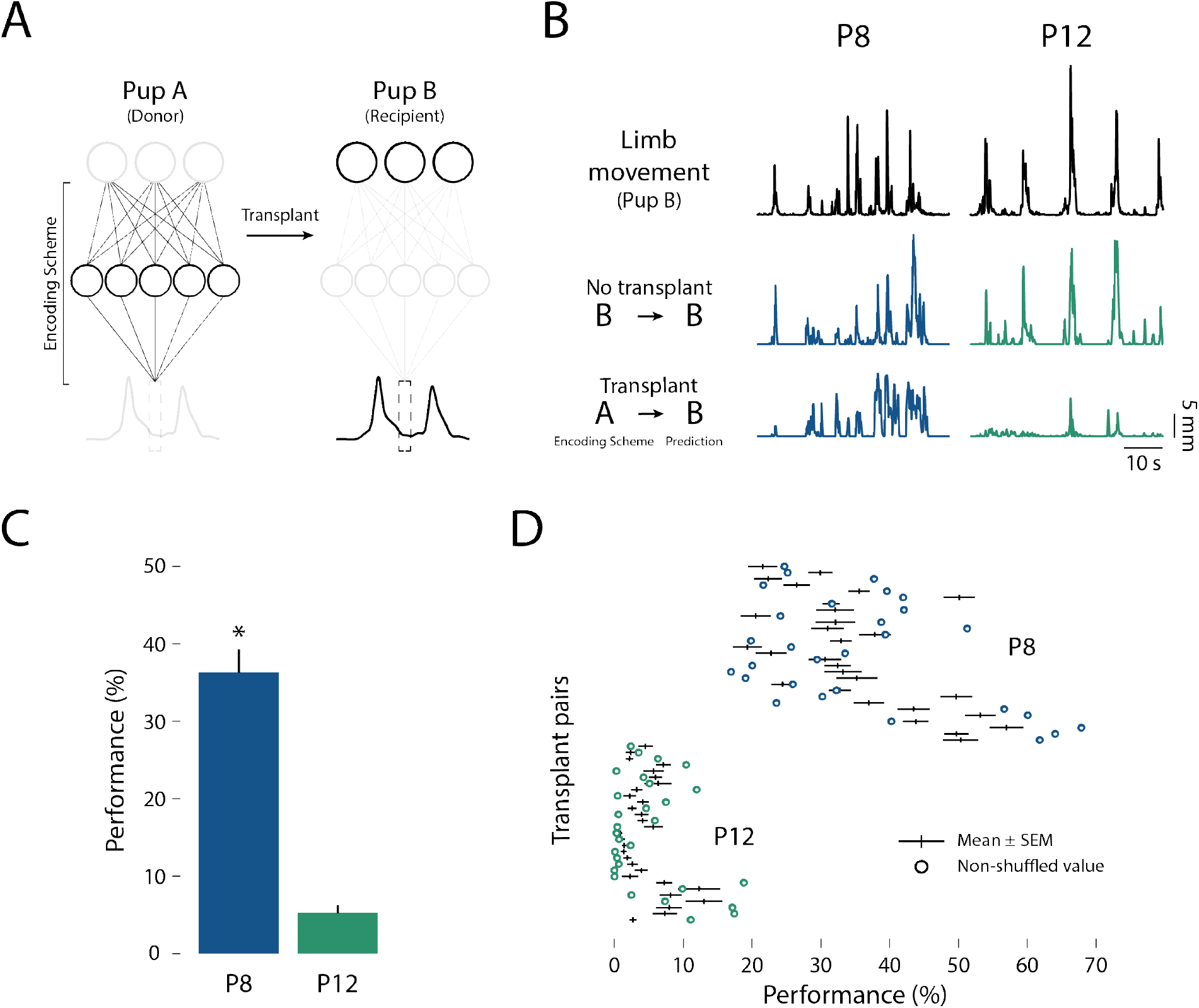
Transplanting M1’s encoding scheme between pups leads to successful decoding at P8, but not at P12. (A)Representation of the “model transplant” experiment. Pup A (“donor”; left) donates its encoding scheme to Pup B (“recipient”; right). Thus, Pup B’s M1 unit activity (top row) is used to predict Pup B’s forelimb displacement (black trace at bottom) using Pup A’s encoding scheme. (**B**) Representative traces of actual limb movement (top) compared to limb movements predicted by Pup B’s (original) decoder model (middle) and limb movement predicted by Pup B’s (transplanted) decoder model (bottom) for P8 (left, blue lines) and P12 (right, green lines) rats. (**C**) Mean (+SEM) decoder performance (measured as the ratio of the “transplant” *r*^2^ to the original *r*^2^, expressed as a percentage) for P8 (blue) and P12 (green) rats. Asterisk denotes significantly better decoder performance after transplantation at P8 than at P12 (p < .05). (**D**) Decoder performance (now on the x-axis) is shown for each donor-recipient pair across 30 random shuffles. Black lines indicate the mean (±SEM) decoder performance of the 30 random shuffles for each donor-recipient pair. Blue and green circles indicate decoder performance of the unshuffled models at P8 and P12, respectively.

To test this prediction, M1 activity of a given P8 or P12 rat (the recipient) was decoded using the model weights (inferred encoding scheme) transplanted from another P8 or P12 rat, respectively (the donor). M1 units were aligned between the donor and recipient rats according to firing rate (i.e., sorted from highest to lowest firing rates). For both P8 and P12 rats, 29 such donor-recipient pairs were generated. (Again, only the nonlinear decoder was used.) As predicted, decoder performance was significantly better across P8 pairings than P12 pairings (t(56) = 11.71, p < .001, adj. *η*^2^ = .705; **Figure 4B–C**). To ensure that this finding was not simply due to our method of aligning firing rates across pups, we performed 30 additional trials of the experiment using random unit-unit pairings; this randomization had no substantive effect on the age-related difference in decoding performance after transplantation (**Figure 4D**). Thus, continuous activity appears to contribute to the emergence of individualized M1 encoding schemes at P12.

## Discussion

We previously reported that M1’s encoding of forelimb movements at P8 disappears with the onset of continuous activity at P12 [6]. That finding raised the possibility that M1’s somatosensory representations are “reset” at P12 by the sudden change in cortical dynamics. An alternative possibility was that the somatosensory representation persists at P12 but is obscured by the noisy continuous activity. We considered the second possibility more parsimonious than the first, in part because twitch-related reafference continues to trigger M1 activity through at least P20 [27]. Accordingly, we predicted that M1 continues to encode movement-related information at P12, but through a more complex encoding scheme. This prediction was confirmed using a nonlinear neural decoder.

We expected the nonlinear decoder to reveal M1 encoding of twitch movements at P12. To our surprise, however, this was not the case; instead, only movement-related information during wake was encoded at P12. Also, wake movements were better reconstructed by the nonlinear decoder at P8, indicating that a significant component of M1’s representation of wake movements is nonlinear even *before* the onset of continuous activity. This last finding is particularly surprising given that the brainstem selectively dampens wake-related reafference in M1 at P8 [20, 21, 31]; moreover, our previous analysis using linear methods led us to conclude that M1 units are not sensitive to wake-related kinematics at either age [6]. Thus, the present finding that M1 encodes twitches and wake movements through linear and nonlinear means, respectively, adds another dimension to our understanding of M1’s state-dependent sensory representation in early development.

It must be stressed that although individual twitches were not reliably decoded at P12, this age does not mark the end of twitches’ significance for sensorimotor development. For example, in P20 rats, twitches are implicated in the developmental emergence of a cerebellar-dependent internal model of movement [27]. To what extent twitching contributes to other forms of plasticity in adults remains an open question [32-34].

### Continuous activity corresponds with more unique M1 information

In adults, continuous activity is associated with complex functions such as sparse coding [28, 35, 36] and predictive coding [35, 37, 38]. Continuous activity is also thought to enhance reafference by providing contextual information related to ongoing behaviors, cognitive processes, and cortical dynamics [39-41]. Accordingly, one might expect the emergence of continuous activity at P12 to immediately sharpen reafference in M1. However, this was not the case (see **Figure 2B**), suggesting that continuous activity *per se* does not enhance reafference and lead to better neural decoding outcomes.

Although continuous activity did not immediately enhance reafference at P12, it did correspond with an increase in the *uniqueness* of information in M1. This unique information could be due, in part, to the developmental narrowing of M1 receptive fields. At P8, M1 units respond across a range of movement amplitudes, suggesting that M1 receptive fields are broadly tuned to multiple forelimb muscles [6]. This broad tuning is no longer apparent at P12, suggesting a progression toward the narrow receptive fields displayed by M1 units in adulthood (Stefanis & Jasper 1964, Nudo et al. 1992, Georgopoulos & Stefanis 2007). Similar receptive-field narrowing has been demonstrated in other cortical (Hubel & Wiesel 1963, DeAngelis et al. 1993) and subcortical (Chen & Regehr 2000, Tschetter et al. 2018) areas and is an activity-dependent process (Chakrabarty & Martin 2005).

The noise that characterizes continuous activity may have a functional benefit related to the concept of “regularization.” Regularization is the process of introducing noise to a system so as to prevent overfitting (i.e., forming inappropriately strong relationships too early in the learning process). Regularization through noise is a conventional technique in the creation of robust computational networks [42-44]. Accordingly, we propose that regularization contributes to receptive-field narrowing by ensuring that connections between M1 units and muscle fibers are only strengthened when they fire together with high-temporal precision exceeding the noise band. Connections with less-precise temporal relations (i.e., at the edges of the receptive field) would be weakened and pruned.

### M1’s encoding scheme individuates at P12

We found that when the encoding scheme of one P8 rat was “transplanted” to another P8 rat, decoding outcomes were significantly better than after a similar “transplant” between P12 rats. In other words, M1’s encoding scheme becomes significantly more individualized, suggesting that the complex encoding schemes at P12 are more specific to individual pups than are the encoding schemes at P8.

Developing animals face the problem of matching cortical connections to a moving target: their rapidly growing bodies. Discontinuous activity solves this problem before P12, as nearly all M1 activity occurs in response to movement-related reafference or external stimulation [6, 8, 45]. This pattern of activity maximizes the correlation between behavior and neural activity. Similarly, before P12, a 10–20 Hz corticothalamic rhythm—known as a *spindle burst*—amplifies reafference and promotes the development of sensorimotor pathways [2, 46-49]. This strong correlation between limb movements and M1 activity is thought to strengthen those M1 connections early in development that will serve as the foundation for later-emerging motor control [6, 21, 45].

## Conclusion

In adults, M1 contributes in many complex ways to motor control and motor learning [50-52]. Also, M1 integrates information arising from the other senses [53, 54], neuromodulatory systems [55, 56], and ongoing behaviors [39-41]. Although little is known about the development of these higher-level functions of M1, it is now clear that these functions—like M1’s most basic motor capacities—rest upon an early-developing sensory foundation. But even this early “sensory phase” of M1 development is protracted and complex: It begins during the discontinuous period of M1 activity as gross somatotopic relations are formed and, as shown here, is transformed through the transition to continuous activity. Thus, the present findings add new dimensions to our growing understanding of M1’s sensory development before it assumes its more familiar role in motor control.

## Acknowledgments

Preparation of this article was made possible by a grant from the National Institute of Child Health and Human Development (R37-HD081168) to M.S.B.

## Author Contributions

Conceptualization: R.G. and M.S.B.; Methodology: R.G.; Software: R.G.; Formal Analysis: R.G.; Investigation: R.G.; Data Curation: R.G; Writing – Original Draft: R.G.; Writing – Review & Editing: R.G., G.S., and M.S.B.; Visualization: R.G. and M.S.B.; Funding Acquisition: G.S. and M.S.B.; Resources: G.S. and M.S.B.; Supervision: M.S.B.

## STAR METHODS

### KEY RESOURCES TABLE

| REGENT or RESOURCE | SOURCE | IDENTIFIER |
| --- | --- | --- |
| **Experimental models: Organisms/strains** |  |  |
| Sprague-Dawley Norway Rats | Envigo | RRID: RGD_10401918 |
| **Software and algorithms** |  |  |
| MATLAB, version 2020b | Mathworks | RRID: SCR_001622 |
| Python Programming Language, version 3.8 | Python Software Foundation | RRID: SCR_008394 |
| Tensorflow, version 2.9 | Google | RRID: SCR_016345 |
| Adobe Illustrator Creative Cloud 2022 | Adobe | RRID: SCR_010279 |
| SPSS 28 | IBM | RRID:SCR_019096 |

### RESOURCE AVAILABILITY

#### Lead contact

Further information and requests for resources should be directed to, and will be fulfilled by, the lead contact, Dr. Mark Blumberg.

#### Materials availability

This study did not generate new unique reagents.

#### Data and code availability

Timestamps of actional potentials, twitches, wake movements, and sleep-wake transitions, as well as the position of the forelimb in Cartesian coordinates are available for download at https://github.com/rglanz/Glanz_et_al_2023. All custom software is available upon request. (Also see https://github.com/KordingLab/Neural_Decoding for neural decoding software and https://github.com/nmtimme/Neuroscience-Information-Theory-Toolbox for information theory software.)

### EXPERIMENTAL MODEL AND SUBJECT DETAILS

All experiments were conducted in accordance with the National Institutes of Health Guide for the Care and Use of Laboratory Animals (NIH Publication No. 80–23) and were approved by the Institutional Animal Care and Use Committee of the University of Iowa (Protocol #: 0021955).

As described previously [6], the data used in this study were collected from Sprague-Dawley Norway rats (*Rattus norvegicus*) at P8–9 (hereafter referred to as P8; n = 8) and P12–13 (hereafter referred to as P12; n = 8). Equal numbers of males and females were used, and all subjects were selected from different litters. See our previous report for further details related to experimental subjects.

### METHOD DETAILS

As described previously [6], high-speed (100 fps) video and M1 extracellular unit activity were collected from unanesthetized pups as they cycled through sleep and wake. Forelimb displacement was quantified using DeepLabCut [57, 58] and forelimb movements were identified using custom software and were visually confirmed. See our previous report for further details related to surgeries, electrophysiological recordings, video data collection, histology, spike sorting, and determination of behavioral state and forelimb movements.

### QUANTIFICATION AND STATISTICAL ANALYSIS

#### Data preparation

For all decoding procedures, M1 unit activity and forelimb displacement served as the predictor and target variables, respectively. Both variables were binned in 20-ms increments. Forelimb displacement was calculated as the absolute value of zero-centered limb position along the x-, y-, and z-dimensions and was smoothed using a 20-ms half-width Gaussian kernel.

To prevent model overfitting, the one-hour recordings were split into training (36 min; 60%) and testing (9 min; 15%) datasets, which were used to train and test model parameters for best performance. A validation dataset (15 min; 25%) was held back until the final model parameters were established. The validation dataset was then used to perform all present analyses in order to ensure an unbiased final assessment of decoder performance [25].

#### Data scaling

M1 activity was z-scored prior to decoding. The scaling factors (mean and standard deviation) were calculated using only the training dataset to avoid data leakage (i.e., positive bias in performance estimates due to information from the testing dataset leaking into the training dataset; [59]. Forelimb position was normalized prior to decoding. The scaling factors (minimum and maximum) were again calculated using only the training dataset to avoid data leakage.

#### Linear model

The linear model was composed of a single-layer Tensorflow model [60] with no hidden layer or activation function. This design forms a linear perceptron [61] and is equivalent to an ordinary least-squares (i.e., linear) regression. All models were assembled in Keras, a software package that assists in building Tensorflow models [62]. A 240-ms time window surrounding each timepoint was flattened (i.e., crossed with the feature of individual units) to provide additional temporal context to the model.

The decoder’s learning parameters were as follows: Mean-square error was selected as the loss function (i.e., a function used to determine the error between the real limb displacement and the predicted limb displacement). Adam [63] was selected as an optimizer (i.e., an algorithm that updates the internal parameters of the model across training iterations). The learning rate (i.e., the degree to which the internal parameters are updated) was set to 0.001 arbitrary units, which is the default learning rate for Adam in the Keras package. Importantly, we tested a variety of different combinations of loss functions, optimizers, and learning rates and did not find meaningful impact on decoder performance (data not shown).

The model was allowed to train until the validation loss function stopped decreasing, which typically occurred after 3–5 iterations. Three target variables were independently predicted for each time bin, corresponding to x-, y-, and z-displacement of the forelimb.

The three predictions were combined using the Pythagorean theorem prior to comparison with actual limb displacement:

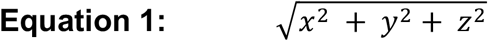

#### Nonlinear model

Several nonlinear models were compared before settling on a long-short-term memory (LSTM) neural network [26]. The LSTM model was chosen for its superior performance on the current dataset and its successful application in similar neural decoding applications [25, 64, 65]. The number of nodes in the input layer was identical to the number of units in each specific M1 recording (i.e., 14–38 nodes). Larger numbers of nodes were tested (up to 500) but did not result in significantly better decoding performance (data not shown). A 240-ms time window, similar to that added to the linear model, was included as a recurrent feature.

To further prevent overfitting, a penalty was added to the recurrent dimension of the model to prevent rapid changes to the model’s internal parameters across training iterations. The penalty selected was L2 (ridge regression; [66]:

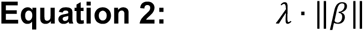

where λ, which represents the magnitude of the penalty, was set to 0.001 arbitrary units. The optimizer, learning rate, and loss function were identical to the linear model. The nonlinear model was also allowed to run until the validation error stopped decreasing (typically within 3–5 iterations). Similar to the linear model, three independent predictions (x-, y-. and z-displacement) were made for each time bin and summed using the Pythagorean theorem (Equation 1) prior to comparison with actual limb displacement.

#### Decoder performance

Decoder performance was evaluated using the square of the Pearson correlation coefficient (*r*^2^) with actual (x) and predicted (y) limb displacement as the two variables:

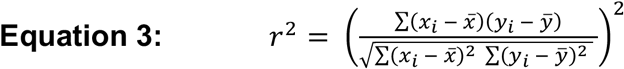

In typical neural decoding applications, regression performance is measured using the Coefficient of Determination (*R*^2^)

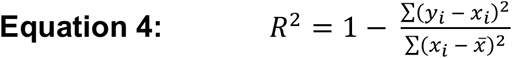

which is sensitive to the scale of the errors. The scale-invariant definition of model performance used here (Equation 3) solves the “intermittent demand” problem [67]. Briefly, because the limb’s displacement is near zero (i.e., at rest) across the vast majority of sampled time points, *R*^2^ disproportionately rewards predictions equal to the mean limb displacement and punishes predictions that vary from baseline, which is not the case with *r*^2^.

#### Lesion experiment

Two “lesion” experiments were performed in which individual M1 units or action potentials (spikes) were randomly selected to be removed from the decoding process. A given percentage of units or spikes, in a range from 0% to 90% (in increments of 10%), was randomly selected and removed without replacement using the NumPy software package in Python [68]. The resulting units or spike trains were then used as the predictor variable and given to the decoder model that was trained on the “intact” dataset. The performance of the decoding model after a lesion was calculated as the ratio of the lesion *r*^2^ to the “intact” *r*^2^, converted to a percentage. The randomization procedure was repeated ten times and the ten *r*^2^ values were averaged (per animal).

#### Mutual Information

Mutual information between a target unit (X0) and a subset of predictor units ({Xk}) of size *k* was defined as:

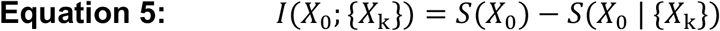

where *I(X*0; {*X*_*k*_}) represents the mutual information between the target unit and subset of units [69-71]. This is equivalent to the entropy of the target unit (*S(X*0)) minus the entropy of the target unit explained by the subset (*S(X*0 | {*X*_*k*_})). Software from [72] was used to calculate mutual information. Random subsets of units were randomly selected from all possible combinations (10 selections per subset of size *k*).

#### Model transplant experiment

To test the interchangeability of decoder models between animals of a particular age, a “model transplant” experiment was performed. The M1 activity (predictor variable) of an individual rat (referred to as Pup A; “recipient”) was used to predict Pup A’s forelimb displacement using the model weights of a different rat (Pup B; “donor”) of the same age. Two animals of the same age were paired when Pup A had the same number of M1 units, or a greater number, than Pup B. The two animals’ units were matched by descending firing rate. The excess units (from Pup A) were discarded from the analysis. In total, 29 such donor-recipient pairs were selected for both P8 and P12 animals. To test whether firing-rate sorting introduced bias, the experiment was repeated 30 times with random unit-unit pairings. The performance of the “model transplant” was measured as the ratio of the transplanted *r*^2^ (Pup A × Pup B) to the original *r*^2^ (Pup A × Pup A).

#### Statistical analyses

Prior to analysis, all data were tested for normality using the Shapiro-Wilk test, for equal variance using Levene’s test (for between-subjects variables), and for sphericity using Mauchly’s test (for within-subjects variables with >2 groups). For analyses in which the variance between groups was not equal, a pooled error term was not used when generating simple main effects and post-hoc tests. For analyses in which sphericity was violated, a Huynh-Feldt correction was applied to the degrees of freedom. All *r*^2^ values were arc-sin transformed prior to analysis. The mean and standard error of the mean (SEM) were used throughout as measures of central tendency and dispersion, respectively.

All analyses were performed as independent t tests or two-way mixed-design ANOVAs. Simple main effects were only tested if the interaction term was significant. In all two-way ANOVAs, an adjusted partial eta-squared was used as an estimate of effect size that corrects for positive bias due to sampling variability [73]. For t tests, an adjusted eta-squared estimate of effect size was reported.

